# Molecular characterization of chikungunya viruses associated with outbreaks in Kenya from 2017 to 2020

**DOI:** 10.64898/2026.02.23.707436

**Authors:** Derrick Amon, Bonventure Juma, Joseph Muriuki, Caroline Ochieng, Naomi Koech, Gilbert Kikwai, Lydia Mwasi, Melvin Ochieng, Caroline Ngugi, Emily H. Davis, Holly R. Hughes, Aaron C. Brault, Naomi Lucchi, Peninah Munyua, Elizabeth Hunsperger

## Abstract

In 2014, Chikungunya virus (CHIKV) infections were reported in Kilifi, Kenya, followed by a major outbreak in Mandera County in 2016. Since then, information regarding subsequent outbreaks in Kenya has been scarce. A study on the burden and etiologies of acute febrile illnesses (AFI) in Kenya reported an increase in CHIKV cases in Mombasa between December 2017 and December 2019. In January and February 2020, another outbreak of CHIKV occurred in Dadaab-Hagadera. These recurrent outbreaks necessitated the establishment of the molecular characteristics and phylogenetic differences of the CHIKVs collected from Mombasa and Dadaab-Hagadera. The challenge of distinguishing chikungunya fever (CHIKF) from other AFIs also underscored the importance of describing predictors of laboratory-confirmed CHIKV cases versus other AFIs. Sequences generated revealed that the Indian Ocean Lineage (IOL) was associated with sporadic CHIKV outbreaks in Kenya from 2017 to 2020. When sequences were compared to the 2014 Kilifi outbreak, key mutations associated with increased CHIKV viral fitness in mosquito vectors, including E1-K211E and E2-V264A, were observed in both Dadaab-Hagadera and Mombasa samples. Time-informed Bayesian phylogenetic analysis demonstrated clear geographic structuring, with Mombasa sequences forming tightly clustered clades and Dadaab-Hagadera sequences grouping separately, indicating sustained local transmission at both sites. However, some conserved amino acid substitutions were shared between the two locations, including nsP1-A104V and E2-I94V, suggesting circulation of closely related lineages. When the association of symptoms to positive CHIKV cases was compared to other AFI symptoms, sore muscles, headache, and convulsions were significantly more common in CHIKV cases. In contrast, diarrhea was considerably less frequent in CHIKV patients. Cough, skin rash, conjunctivitis, and vomiting were not significantly associated with CHIKV infection. In regions with limited access to point-of-care diagnostics, identifying clinical symptoms that are likely to be strongly associated with CHIKV infection can be important in the differential diagnosis from other AFIs, facilitating reliable patient screening and improving diagnostic accuracy, particularly during outbreaks when timely identification of CHIKV is critical. Continuous molecular surveillance of circulating CHIKV genotypes is important as the mutations identified in these CHIKV strains continuously accumulate and may have a significant impact on control strategies.

## Introduction

Chikungunya virus (CHIKV) is a vector-borne alphavirus in the family *Togaviridae* and is associated with severe morbidity in affected populations, including acute fever and polyarthritis [1]. First identified in Tanzania in 1952[2], CHIKV has subsequently been associated with disease outbreaks and is a notable cause of re-emerging infectious diseases with global outbreaks reported in Asia, Latin America, the Pacific Islands, and Africa. CHIKV was initially considered to be non-life-threatening, but reports of an outbreak in La Réunion in 2006 revealed clinical manifestations of neurological and hemorrhagic symptoms that were also linked to mother-to-child transmissions[3]. Symptomatic CHIKV attack rate among infected persons can be as high as 72%[4]. In Kenya, CHIKV is endemic in the coastal region, with a high burden in children below one year[5]. In December 2018, data revealed that the annual incidence of CHIKV-associated neurological disease in coastal Kenya varied from 13 to 58 episodes per 100,000 person-years in children below 16 years old, a clear indication that CHIKV-associated neurological disease is common and should be considered a public health concern[6]. Coastal Kenya has experienced recurrent outbreaks of both dengue and CHIKV over time and continues to be at risk for other acute febrile illnesses (AFIs), including malaria [7]. During such outbreaks, CHIKV infections can be easily mistaken for other AFIs, as they share similar symptoms that are difficult to distinguish, particularly in areas with limited laboratory capacity for confirmation. This can lead to misdiagnosis and underreporting of CHIKV cases, eventually straining public health systems.

The first major CHIKV outbreak in Kenya occurred in 2004 in Lamu, with an estimated 13,500 cases detected[8]. This outbreak spread to the coastal city of Mombasa and the Indian Ocean islands of Comoros and La Réunion, with over 500,000 cases reported [9]. In 2014, CHIKV cases were detected in Kilifi County along the northern coast of Kenya [10]. This was followed by another outbreak in 2016, where CHIKV cases were confirmed in the Bula Hawa region of Somalia, with the outbreak eventually spreading to Mandera County, Kenya [11]. Joint efforts between Kenya and Somalia helped contain the outbreak, but this marked a resurgence of CHIKV activity in Kenya. Between 2017 and 2019, CHIKV cases were reported among patients presenting with fever at Coast General Hospital in Mombasa, according to a surveillance study on acute febrile illnesses[7]. In January 2020, another outbreak occurred in the Dadaab Refugee Camp, located in Garissa County, which is also near the Kenya-Somalia border. The camp comprises of 3 camps, Hagadera, Dagahaley, and Ifo. It hosted approximately 385,328 refugees as of July 2024[12], and continues to grow due to ongoing cross-border movement, making it highly susceptible to the introduction and sustained circulation of CHIKV and other arboviruses. Apart from northern and coastal Kenya, other regions have recorded very few cases of CHIKV, but laboratory studies have demonstrated that *Aedes aegypti* populations from areas like Nairobi and Kisumu are all capable of transmitting CHIKV[13]

Phylogenetic analysis of CHIKV using the E1 gene sequence defines four distinct genotypes[14] The East-Central-South African (ECSA) and the West African genotypes are endemic in sub-Saharan Africa and associated with epidemics within this region. The Asian genotype and the Indian Ocean lineage (IOL) are associated with epidemics in parts of Asia and the Indian Ocean islands, respectively, with the IOL having diverged from the ECSA genotype[15].

*Ae. aegypti* is the principal vector for CHIKV transmission; however, *Aedes albopictus* subsequently emerged as the primary vector in the reported outbreaks in La Réunion Island, Europe, and Gabon[16][17]. The La Réunion outbreak was marked by mutations that enhanced the viral fitness of CHIKV in *Ae. albopictus[18]*, likely contributing to the wide geographical spread of CHIKV during this period. Enhanced viral fitness of CHIKV was linked to the E1:K211E and E2:V264A mutations, which have also been associated with increased dissemination and transmission in *Ae. aegypti* [11], thus posing a risk of extensive outbreaks in previously unaffected areas. CHIKV structural glycoproteins E1, E2, and E3 play a crucial role in the interaction of the virus with host cells, impacting both the efficiency of transmission and the host’s immune response. Glycoprotein E1 is responsible for mediating cell fusion, while E2 facilitates interaction with host cell receptors[19]. E3 assists in the heterodimerization of E1 and p62, preventing premature exposure of the E1 fusion loops[19]. Adaptive mutations, particularly in these envelope proteins, may enhance the virus’s ability to circulate and persist in endemic regions, potentially heightening the risk of more extensive and severe CHIKV outbreaks[19]. In light of these reported outbreaks in Dadaab-Hagadera and Mombasa, we characterized viral genomes collected from these two regions using whole-genome sequencing and compared them with genomes from previous outbreaks. We also analyzed CHIKV-associated symptoms that could help in differential diagnosis during such outbreaks.

## Methods

### Sample Collection

This was a retrospective study of CHIKV-positive samples archived at the Kenya Medical Research Institute-Centre for Global Health Research (KEMRI-CGHR) Laboratory, supported by the U.S. Centers for Disease Control and Prevention (US CDC) from December 2017 to February 2020. Out of the 42 samples analyzed, 32 were from an acute febrile illness (AFI) surveillance in which whole blood and serum samples were collected from patients at the Coast General Hospital in Mombasa and tested for a wide range of targets using an AFI TaqMan Array Card (TAC, Applied Biosystems, USA)[7]. Nine samples were collected from an outbreak in Dadaab-Hagadera in 2020 and were tested using CHIKV RT-PCR assay (CDC, Fort Collins, CO, USA). An additional sample was added from a virus isolated from one of the serum samples collected in Dadaab-Hagadera and passaged three times in Vero cells.

For AFI cases, paired whole blood and serum samples were collected from the same participant. Whole blood was used for initial TaqMan Array Card TAC testing, and corresponding serum aliquots were immediately archived at −80 °C. Total nucleic acid extracted from whole blood was also stored at −80 °C after TAC testing. For outbreak investigation samples, in which only serum was collected, serum specimens were archived at −80 °C after initial laboratory analysis. AFI cases were defined as inpatients with a temperature ≥38.0°C on admission and <14 days onset prior to presenting at the facility[7]. For cases of undifferentiated fever (UF), defined as AFI without diarrhea (3 loose stools in 24 hours) or lower respiratory tract symptoms (cough/difficulty breathing plus oxygen saturation <90% or [in children <5 years] chest indrawing) whole blood samples were collected and tested using a real-time PCR-based TAC for 17 viral, 8 bacterial, and 3 protozoal pathogens. After obtaining participants’ consent, trained surveillance officers conducted interviews using a standardized questionnaire to collect information on demographic, socioeconomic, clinical, and risk factors. For participants younger than 18 years, consent was provided by a parent or legal guardian, with additional assent obtained from children aged 7–17 years. Consent and assent forms were read aloud to participants in a language they understood. Written consent was obtained from literate participants, whereas witnessed consent was used for those unable to read or write. For children younger than 7 years and for individuals lacking decision-making capacity due to severe illness or cognitive impairment, consent was obtained solely from a parent or legal guardian. This was followed by a physical examination where clinical symptoms, including headache and myalgia, were ascertained through clinician assessment and caregiver report for younger children, as part of standardized surveillance questionnaires. To ensure quality, all medical charts were reviewed once the enrolled patients were discharged, and data on clinical course, management, and outcome were abstracted[7]. Samples were shipped to the KEMRI-CGHR laboratory in Nairobi within 7 days. For Dadaab-Hagadera, serum samples were collected following a rise in documented febrile illness cases in the region. The Kenya Ministry of Health collected specimens according to their established protocols from January 2020. These samples were transported to the KEMRI-CGHR lab for CHIKV testing. The two study locations (Figure 1), which also shows the distribution of CHIKV cases in different districts in Mombasa.

**Figure 1.**
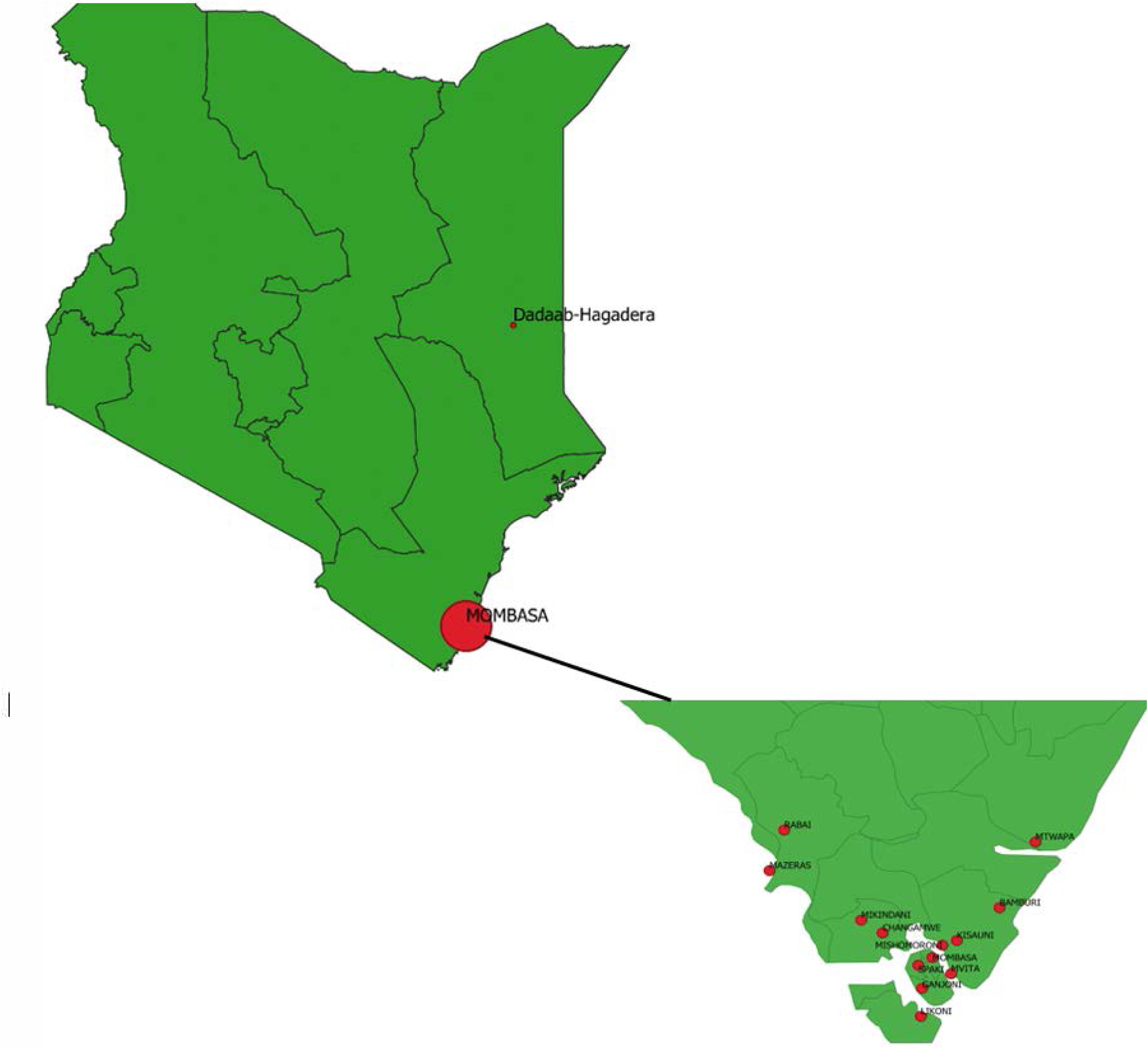
Map of Kenya showing the two region affected by CHIKV outbre

### Nucleic Acid Extraction and qRT-PCR

Archived whole blood samples were used to extract total nucleic acid (TNA) using Highpure high volume TNA Extraction Kit (Roche, USA) following the manufacturer’s instructions and eluted in 150 µl elution buffer. Bacteriophage MS2 and Phocine Herpesvirus (PhHV) were added to each sample as extrinsic controls to validate the extraction process and confirm that total nucleic acids (both DNA and RNA) were amplified successfully. For the Dadaab-Hagadera outbreak, serum samples were extracted using a Qiamp Viral RNA isolation kit (Qiagen Inc, Valencia, CA).

Samples collected for AFI surveillance from the Coast General Hospital were tested using the TaqMan Array Card (TAC) designed to test multiple pathogens, including CHIKV[7]. 46 µl of the TNA was mixed with 54 µl AgPath One Step RT-PCR master mix buffer (Life Technologies Corporation). The 100 µl total volume was pipetted into the loading port of the AFI TAC card and the cards were centrifuged at 15294 X G for 1 minute twice. The cards were sealed using the Taqman Low-density array (TLDA) sealer and the loading ports were removed. The TAC card assay was performed on the ViiA™ 7 real-time PCR system (Life Technologies Corporation, USA) platform using PCR cyclic conditions of 20 minutes at 45 °C, 10 minutes at 95 °C followed by 45 two-step cycles of 15 Seconds at 95 °C and 1 minute at 60 °C. Samples were designated positive if the cycle threshold (Ct) value for a target was ≤ 37.

Outbreak samples from Dadaab-Hagadera were tested for CHIKV using the CHIKV-specific, real-time RT-PCR following the manufacturer’s instructions with thermocycling conditions similar to the TAC Assay conditions using published primer-probe set targeting the CHIKV non-structural protein 4 (nsP4) region (Table 1) [20]

### Viral Isolation and extraction from viral isolates

Forty-one human sera samples archived after testing positive for CHIKV RNA were subsequently utilized for virus isolation. Minimum Essential Medium (MEM) (Gibco, Thermo Fisher Scientific, Waltham, MA, USA). containing 10% fetal bovine serum, 2% L-glutamine, and 1% antibiotics in T-25 tissue culture flasks were used to grow Vero cells to about 95% confluency. 100 µl serum was mixed with 300 µl MEM and inoculated onto Vero cells[21].

Inoculated cells were incubated at 37 °C with 5% CO_2_ for 12 days. Mock-inoculated Vero cell controls containing culture medium only were included and maintained in parallel with inoculated cultures. These controls were monitored throughout the incubation period, and no cytopathic effects were observed. Cell cultures were maintained under appropriate supplemented conditions, and sodium bicarbonate was supplemented as needed to prevent buffering depletion. The development of the cytopathic effect was monitored, and once observed in a majority of the cells, the flask was frozen at −80 °C and freeze-thawed to facilitate cell lysis and virus release[22]. For removal of cell debris, centrifugation of the isolates was done at 956 Relative Centrifugal Force (RCF) for 10 minutes. The supernatant containing the virus was transferred into cryovials and stored at −80 °C.

Viral RNA was extracted from isolates using the Qiamp Viral RNA isolation kit (Qiagen Inc, Valencia, CA) following the manufacturer’s protocol and confirmed using CHIKV RT-PCR through the Agpath-ID^TM^ One step RT_PCR Kit (Life Technologies, CA, USA).

### CHIKV Sequencing

Primer sets (Appendix 1) were designed using primal scheme V 1.3.2 to cover at least 99% of the CHIKV genome[23]. The primers were based on references archived in the National Center for Biotechnology Information (NCBI). The primer sequences were manufactured in CDC Atlanta and validated to ensure they are specific and have high annealing efficiency with the right concentration by testing different concentrations using the Agilent 2100 Bioanalyser. Upon validation, primers were shown to have good specificity and amplification efficiency with a concentration of 20 µM.

RNA from one successfully harvested isolate, thirty-two archived total nucleic acids (TNA) from AFI whole blood samples, and nine RNA extracted from serum samples from Dadaab-Hagadera were used for sequencing. cDNA synthesis and library preparation were done using the Illumina COVIDSeq (Illumina, San Diego, CA, USA) kit, following the manufacturer’s instructions and substituting the COVID primers with study-specific CHIKV primers in Appendix 1. Libraries were pooled and cleaned to remove fragment sizes that were either too large or too small by binding the optimal-sized fragments to magnetic beads. The library concentration was determined using a Qubit dsDNA HS Assay Kit (ThermoFisher Scientific Inc., Wilmington, DE, USA) and normalized to 4nM. Sequencing was done through an Illumina Miseq platform using the Miseq V2 300 cycle reagent kit (Illumina, San Diego, CA, USA). This generated base paired-end reads. Read quality was determined using the FASTQC tool and low-quality reads and primer dimers were trimmed off to ensure good-quality reads were passed on to downstream analysis.

### Data analysis

After quality analysis, read alignment and mapping of sequenced reads to reference genomes were done using Bowtie2 (V2.2.5)[24]. LoFreq (V2.1.5) was used for variant detection, enabling the identification of single nucleotide variants (SNVs) with the CHIKV viral genome[25]. For de novo assembly, SPAdes (V1.2.7)[26] was used, employing a multi-k-mer approach to construct contigs from the aligned reads. To refine the assembly and enhance accuracy, Trimmomatic (V0.22) was used for trimming low-quality bases[27], and SAMtools (V2.1.2) for manipulating sequence alignment data[28]. Picard tools were used to mark and remove PCR duplicates from the dataset. Bayesian phylogeny was inferred in Beast v 1.8.4using the TN92+G nucleotide substitution model, an uncorrelated relaxed clock with 100,000,000 MCMC states and 10% burn-in. A substitution model test was performed in MEGA (V7.0.18)[29]. The samples were compared to four different CHIKV lineages obtained from the National Center for Biotechnology Information (NCBI) database: East African (HQ456253.1 CHIKV strain Com125) ECSA Novel (MN402891.1 CHIKV isolate 19CHKYN07LHX) West African (KP003812.1 CHIKV strain GABOPY1) and Indian Ocean Lineage (Kenyan MT526802.1 CHIKV isolate KLF_69219) that caused the 2014 outbreak in Kilifi, Kenya. A total of 50 consensus genomes were included to determine the phylogenetic placement of the Mombasa and Dadaab–Hagadera sequences. Representative genomes were selected to minimize redundancy and simplify phylogenetic inference.

The clinical data from patients with confirmed CHIKV infection (n = 31) and patients with other acute febrile illnesses (AFIs) (n = 741) were analyzed to identify clinical predictors associated with CHIKV. The analysis focused on comparing the presence of symptoms between CHIKV and other AFI cases. The Chi-square test was used to assess associations between symptoms and CHIKV when all expected cell counts were greater than or equal to 5. If any expected cell count was below 5, Fisher’s exact test was applied [30]. p-values were reported alongside (odds ratio) ORs and (confidence interval) CIs, with statistical significance defined as p<0.05. All analyses were conducted in R statistical software (version 4.4.2) (R Foundation for Statistical Computing, Vienna, Austria).

### Ethics Statement

This study was approved by the Kenya Medical Research Institute Scientific and Ethical Review Unit under Study Number (SSC 4116). The study was also approved by the Kenya National Commission for Science and Technology and Innovation (NACOSTI) with license number NACOSTI/P/22/15382. The Acute febrile illness surveillance study was approved by the Kenya Medical Research Institute’s Scientific and Ethical Review Unit (Number SSC 2980) and the Institutional Review Board for the US Centers for Disease Control and Prevention (protocol number 6757).

## Results

### Clinical characteristics of CHIKV cases

In total, 31 out of the 32 confirmed CHIKV cases that had complete datasets and 741 other AFI cases were used to describe clinical characteristics of CHIKV cases compared to other AFIs (Tables 2 and 3). Sore muscle was significantly more common in CHIKV cases (10% vs. 0%, p < 0.001), as were headache (6% vs. 0%, p = 0.0016) and convulsions (61% vs. 18%, p < 0.001). In contrast, diarrhea was significantly less frequent in CHIKV cases compared to other AFIs (3% vs. 32%, p = 0.0013), suggesting it may be a negative predictor of CHIKV. Other symptoms, including cough (43% vs. 40%, p = 0.283), skin rash (3% vs. 6.6%, p = 0.7053), conjunctivitis (5% vs. 0.2%, p = 0.1158), and vomiting (26% vs. 27%, p = 1.000), were not significantly associated with CHIKV. The absence of headache in the AFI group may reflect underreporting, particularly given the low median age of 2.5 years in this group, where younger children are unable to reliably communicate headache symptoms. However, these results suggest that sore muscles, headache, and convulsions are strong clinical indicators of CHIKV, while diarrhea appears to be less prevalent in CHIKV cases. Samples collected from Dadaab-Hagadera had limited clinical information and were not included in the analysis of clinical characteristics.

### CHIKV Isolation

Out of the 41 sera used for viral isolation, CHIKV was successfully isolated from one sample (S37) collected in Dadaab-Hagadera. Following three passages, the isolate (S32) was harvested and included in the sequencing analysis. The isolate was assigned a separate identifier (S32) during sequencing for laboratory tracking purposes.

### CHIKV Sequencing

A total of 42 samples were sequenced. This included archived TNAs from AFI surveillance (32), extracted RNA from archived Dadaab-Hagadera serum samples (9) and the (1) positive virus isolate. 4 samples had incomplete sequence data and were therefore excluded from analysis. The remaining 38 had excellent coverage of 95% and above. Assembled sequences were deposited in GISAID EpiArboTM with the virus names and accession numbers in Appendix 2.

Both the Mombasa and Dadaab-Hagadera strains clustered within CHIKV Indian Ocean sub-lineage (IOL) ECSA (East/Central/South African) (Fig 2). A phylogenetic tree was constructed using sequences of the four lineages obtained from NCBI, which showed that most samples clustered within the location collected. Strains from coastal Kenya, including Mombasa, Kilifi, Changamwe, Bamburi, Mishomoroni, Likoni, and Mtwapa districts, formed a clustered clade characterized by short branch lengths, a sign of sustained local transmission during the 2017–2018 coastal outbreaks. This was also observed in the Dadaab-Hagadera strains, which formed a distinct clade separate from the Mombasa samples (Fig 3).

**Figure 2.**
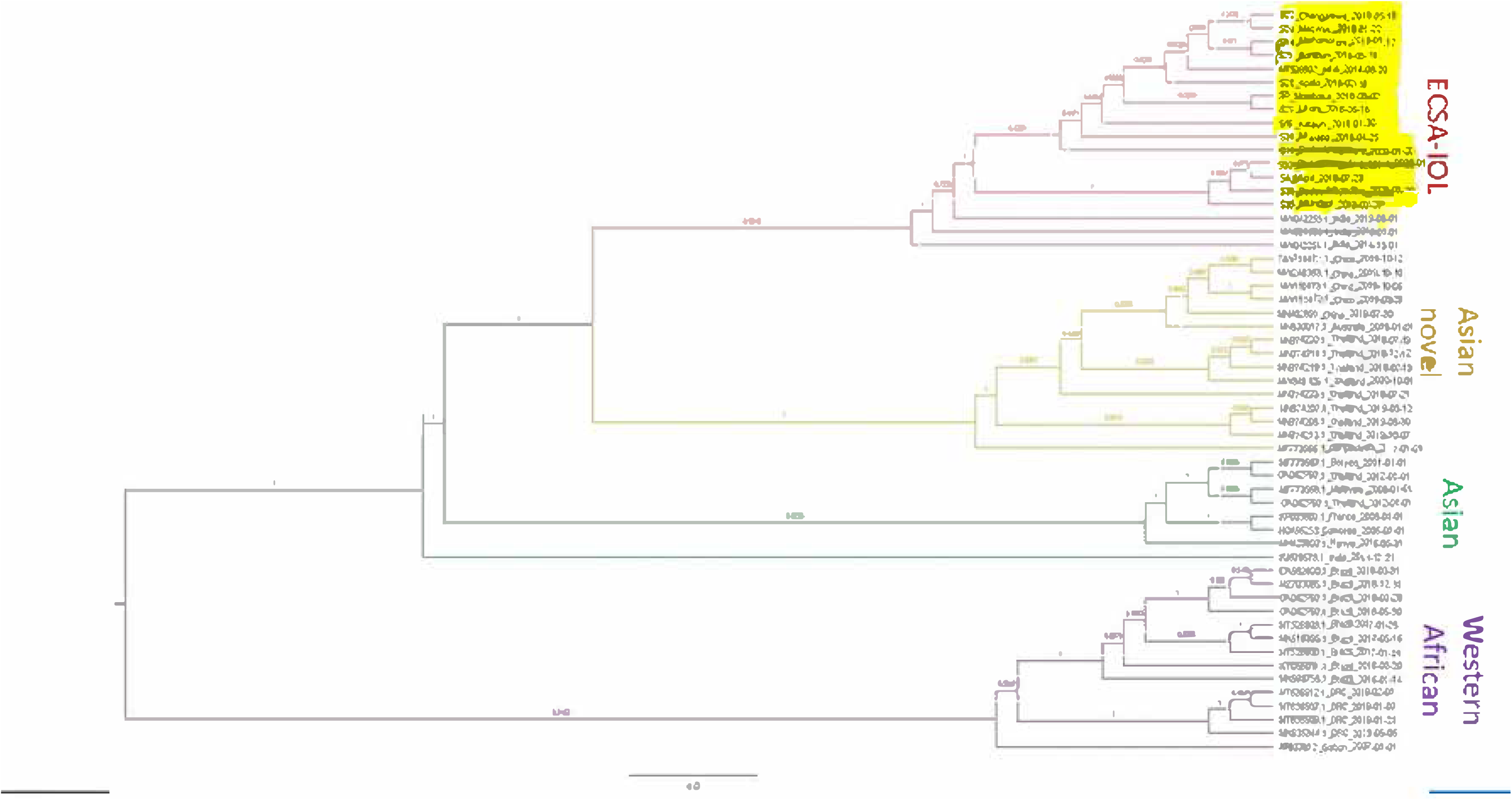
Phylogenetic tree generated using 100,000,000 states and 10% b

**Figure 3.**
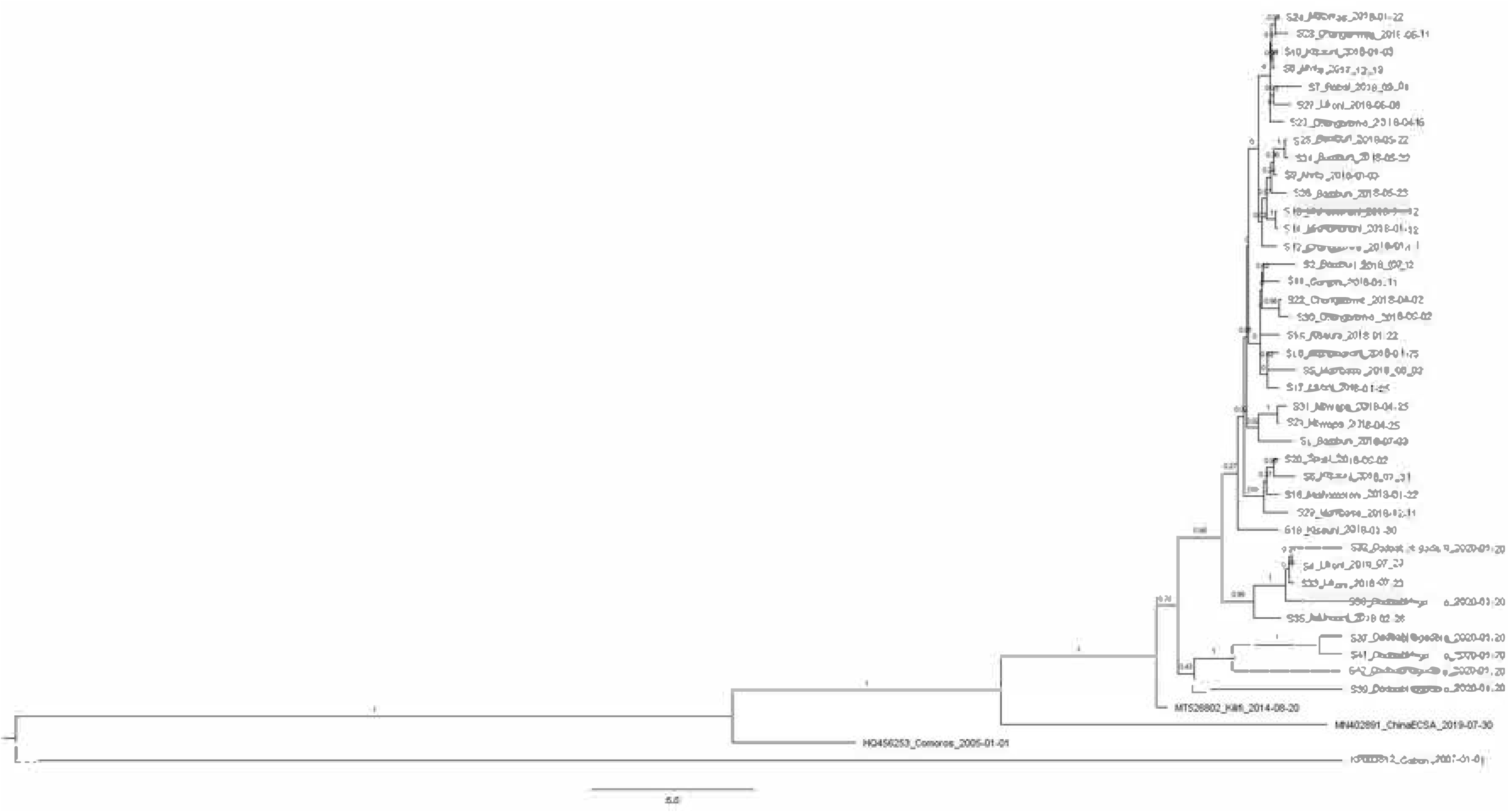
CHIKV from coastal Kenya, forming a tightly clustered clade char

Comparison of consensus sequences for sample S37, which was passaged three times in Vero cells, yielding isolate S32, identified non-synonymous substitutions in nsP1 and E2.

Compared to Dadaab-Hagadera samples, Mombasa samples were more distinct when compared to the reference strain, Kenyan isolate in the IOL sub lineage (MT526802.1 CHIKV isolate KLF_69219). Analysis of single-nucleotide variants (SNVs) revealed clear differences in mutation profiles between the Mombasa and Dadaab–Hagadera CHIKV populations. Among the 32 Mombasa genomes, recurrent amino acid substitutions were predominantly observed in the envelope glycoproteins and non-structural proteins. The E1:A226V mutation was detected in 100% (32/32) of Mombasa samples, consistent with circulation of the Indian Ocean Lineage (IOL). Additional E1 substitutions included E1:T82I, present in 81.3% (26/32) of samples (S1, S2, S4–S15, S17–S25, S27, S28), and E1:V84D, detected in 62.5% (20/32) (S5–S12, S14–S21, S23, S24). The E2:I94V substitution occurred in 68.8% (22/32) of Mombasa samples, while nsP1:A104V was observed in 75.0% (24/32) (Table 4). The Dadaab–Hagadera sequences formed a genetically distinct profile. All samples shared the E1:A226V substitution (100%, 5/5), but lacked several mutations frequently observed in the Mombasa cohort. The E2:I94V mutation was detected in 20% (1/5) (S37 only), while E1:T82I and E1:V84D were absent (Table 4). Substitutions in non-structural proteins were limited, with nsP2 and nsP3 mutations appearing sporadically, each observed in ≤40% of samples and without consistent recurrence across the dataset. Common conserved changes between Mombasa samples and Dadaab-Hagadera samples included nsP1-A104V and E2-I94V (Table 4). Another change between all Mombasa samples and the and the 2014 Kilifi CHIKV strain was located at nsP3-L290F. Other additional changes included sample S43, nsP4-T258A, and sample S37, E2-H227R. All samples contained E1:A226V and E1-K211E/E2-V264A substitutions, which have been associated with increased transmissibility in *Ae. aegypti [31]*.

## Discussion

CHIKV remains a significant emerging public health challenge globally, due to the recurring nature of its outbreaks. In Kenya, since the first major CHIKV outbreak in 2004 along the Kenyan coast, sporadic cases have been reported[8]. In this study, direct sequencing of CHIKV genomes was performed using clinical samples collected in Kenya between December 2017 and February 2020 in two different regions: Mombasa in coastal Kenya, and Dadaab-Hagadera, which is a major refugee site close to the Kenya-Somalia border. Our findings confirm that the mutations associated with increased viral fitness found in previous outbreaks in India, Pakistan, and the 2016 outbreak in Mandera[11], were also present in the 2017–2020 outbreaks in Kenya. *Aedes aegypti* is the predominant vector responsible for transmitting CHIKV in Kenya, and one of the fitness-enhancing mutations, E1-K211E, was detected in the CHIKV samples analyzed in this study [13]. This mutation, along with the E1-A226V and the E2:V264A mutation, had previously contributed to large outbreaks in other regions[32]. The high prevalence of E1:T82I (81.3%) and E1:V84D (62.5%) in Mombasa strains is notable. These substitutions lie within the CHIKV fusion loop and are adjacent to previously described adaptive sites, including V80A [19]. Studies have demonstrated that substitutions at position E1-82 may arise as secondary adaptive steps that may potentially enhance viral fitness [19]. The presence of shared mutations such as nsP1-A104V and E2-I94V in both Mombasa and Dadaab-Hagadera samples suggests possible circulation of genetically linked strains and could also be an indication of convergent evolution driven by ecological or vector pressures. The nsP3-L290F mutation, being observed exclusively in Mombasa samples, suggests region-specific viral adaptation, possibly influenced by local vector population and environmental factors. Further research is necessary to assess whether these additional mutations could give rise to new CHIKV variants that may outcompete and replace the previously circulating strains, altering transmission dynamics and leading to larger and more sustained outbreaks.

Phylogenetic analysis revealed that CHIKV strains collected in Mombasa and Dadaab-Hagadera belonged to the Indian Ocean lineage, supporting previous reports[18]. IOL is known for its increased viral fitness to *Ae. albopictus* due to the E1-A226V mutation. As is common in IOL, all the samples analyzed in this study contained this mutation. Although this mutation is well established, continued detection alongside other mutations suggest ongoing selective pressures that could enhance CHIKV transmission, posing a risk of persistence in new ecological niches and likelihood of larger outbreaks. The genetic variability observed among CHIKV lineages, as evidenced by the distinct consensus level changes between the Kenya outbreak samples from 2017 to 2020 and the Kenyan reference isolate (MT526802.1), underscores the dynamic nature of CHIKV evolution. These findings align with previous studies highlighting the adaptive capacity of CHIKV in response to selective pressures[33].

Viral passages in cell cultures are commonly associated with the accumulation of positively charged amino acid changes in structural genes[34]. This has been known to assist in binding glycosaminoglycans (GAGs) and allow the virus to attach to cells more efficiently. Sample S37 was successfully passaged three times, giving rise to sample S32. Similar to findings in previous studies[35], this passage likely induced non-synonymous mutations in key viral genes, including nsP1 and E2, leading to amino acid changes that may enhance the virus’s ability to attach to host cells more efficiently. The nsP1 gene plays a critical role in viral replication and host interaction, and mutations in this region could improve the virus’s ability to hijack the host immune response[36]. The accumulation of glycoprotein and non-structural protein substitutions in Mombasa strains raises concern about the potential for viruses with enhanced capacity for efficient transmission. The low viral isolation rate may be attributed to sample degradation, as the samples were retrieved for sequencing after being stored for more than three years.

Differentiating the clinical features of confirmed CHIKV cases from other AFIs can be particularly valuable during outbreaks, especially in settings lacking point-of-care diagnostics. Rash and joint pain have been identified as the most effective positive predictors of CHIKV in other regions[30], but data on these clinical features in Kenya has been limited. In this study, we identified key distinguishing characteristics of CHIKV infections based on outbreaks along the Kenyan coast. Sore muscles, headache, and convulsions were more strongly associated with CHIKV patients than in those with other AFIs. In very young children, particularly infants and toddlers, headache is difficult to assess or communicate, likely leading to underreporting or misreporting. This highlights an important limitation in symptom-based diagnosis between CHIKV and other AFIs, underscoring the need for laboratory confirmation. Diarrhea was significantly less common in CHIKV patients compared to those with other AFIs as reported in previous studies, indicating that gastrointestinal symptoms are not prominent features of CHIKV infections[37]. Diarrhea can therefore be a useful criterion to rule out CHIKV infection when present in a clinical setting. Symptoms such as cough, skin rash, conjunctivitis, and vomiting were not significantly associated with CHIKV. Information on joint pain, a common symptom of CHIKV infection, was not collected in this study and could not be assessed as a potential clinical predictor. The young age of the CHIKV group could have led to overrepresentation of some of the symptoms, like convulsions, which were unexpectedly high, necessitating further investigations into age-specific clinical presentations. This study was also limited by a small size of less than 50 confirmed RT-PCR cases hence the need for more studies distinguishing CHIKV clinical features from other AFIs.

## Conclusion

The molecular characterization performed for CHIKV in this study identified ECSA-IOL as the genotype that caused sporadic outbreaks in Kenya from 2017 to 2020. Subsequent to our data sampling period, CHIKV outbreaks were reported in Kenya in 2022[38] underscoring the virus’s continued presence and the need for continuous molecular surveillance that can help improve strategies for control and prevention of future outbreaks. The information provided on the diversity of CHIKV strains can improve the range of understanding of the viral proteins that can be targeted by vaccines. More studies are needed to establish specific signs and symptoms of CHIKV compared to other AFIs in the Kenyan setting for timely diagnosis and management during outbreaks.

## Authorship disclaimer

The findings and conclusions in this report are those of the authors and do not necessarily represent the official position of the funding agencies. Use of trade names is for identification only and does not imply endorsement by [the Centers for Disease Control and Prevention/the Agency for Toxic Substances and Disease Registry], the Public Health Service, or the U.S. Department of Health and Human Services.”

## Acknowledgments

We gratefully acknowledge the surveillance officers and staff of the Kenya Ministry of Health for their efforts in identifying CHIKV cases at Mombasa County Referral Hospital and Dadaab–Hagadera. We extend our appreciation to Godfrey Binaisa (KEMRI-CGHR) for retrieving the samples from storage for processing. We are also grateful to the participants of the AFI study for their willingness to provide information and biological samples despite being acutely ill. We also thank the Division of Vector-Borne Diseases, Arboviral Diseases Branch in Fort Collins, Colorado, U.S.A., for their valuable assistance with the bioinformatics analyses.

## Author Contributions

**Conceptualization:** D. Amon, E.A. Hunsperger.

**Data curation:** D. Amon, M. Ochieng, C. Ochieng, N. Koech, G. Kikwai, L. Mwasi, E.H. Davis, H.R. Hughes, A.C. Brault.

**Formal analysis:** D. Amon, E.H. Davis, H.R. Hughes, A.C. Brault, E.A. Hunsperger.

**Funding acquisition:** E.A. Hunsperger, P. Munyua, B. Juma.

**Investigation:** D. Amon, E.A. Hunsperger, C. Ngugi, J. Muriuki, N. Lucchi, P. Munyua, B. Juma.

**Methodology:** D. Amon, E.A. Hunsperger, P. Munyua, B. Juma, N. Lucchi, C. Ochieng, N. Koech, G. Kikwai, L. Mwasi, E.H. Davis, H.R. Hughes, A.C. Brault.

**Project administration:** D. Amon, E.A. Hunsperger, C. Ngugi, J. Muriuki, N. Lucchi, P. Munyua, B. Juma, E.H. Davis, H.R. Hughes, A.C. Brault.

**Resources:** D. Amon, E.A. Hunsperger, N. Lucchi, B. Juma.

